# The Neural Basis of Habit Formation Measured in Goal-Directed Response Switching

**DOI:** 10.1101/2025.03.13.643040

**Authors:** Mario Michiels, Vincent Man, David Luque, Ignacio Obeso

**Affiliations:** CINAC, Hospital Universitario HM Puerta del Sur, Madrid, Spain; Division of the Humanities and Social Sciences, California Institute of Technology, Pasadena, CA, USA; Departamento de Psicología Básica, Universidad de Málaga, Málaga, Spain; Instituto de Investigación Biomédica de Málaga y Plataforma en Nanomedicina (IBIMA-Plataforma BIONAND), Málaga, Spain; Cajal Neuroscience Centre, Consejo Superior de Investigaciones Científicas (CSIC), Madrid, Spain

**Keywords:** habits, premotor cortex, computational modeling, fMRI, TMS

## Abstract

To override ongoing habitual responses requires switching well-learned actions with new goal-directed processing. However, the neural circuits responsible for these processes remain unclear. This study infers habit strength by introducing a novel task capturing the increased cost associated with switching a habitual response. We employed neuroimaging and brain stimulation to examine the dynamic interactions between human brain regions involved in habits and their interference with ongoing incompatible goal-directed behavior. Training S-R links in overtrained stimuli (compared to less trained ones, termed standard-trained stimuli) increased RT switch costs, explained by drift diffusion computations governing both the training and outcome devaluation phases. Training engaged sensorimotor areas and the posterior putamen, whereas standard trained behaviors, recruited the posterior caudate, insula, and prefrontal regions. A cortical network orchestrated habit expression (right S1 with the left anterior insula/prefrontal areas) while also implicating basal ganglia when overriding habits (left premotor with the putamen). Importantly, stimulation of the left premotor played a causal role in habit control, enhancing performance across both the training and devaluation phases. Our findings reveal an interaction between habitual and goal-directed brain regions, highlighting shared neural dynamics when overriding habitual behaviors.

## Introduction

Changing established habits is often necessary. Successfully overriding an old habit requires considerable effort, and initially, greater mental costs arise due to the intrusion of the acquired habit. The additional cost of overriding habits reflects the conflict between two distinct, competing systems (Balleine and O’Doherty 2010). The first is the goal-directed system, in which actions are executed by prospectively evaluating the behavioral options to achieve task goals, considering the available knowledge of the task and environmental context. Goal-directed actions are, therefore, computationally demanding and slow. The second is the habit system, where stimulus-driven responses are executed automatically. These responses are gradually acquired through repeated practice (i.e., by trial and error) and then applied in a highly efficient manner, enabling very rapid execution. However, this efficiency comes at the cost of inflexibility, making habitual behaviors difficult to override.

Recent habit research in humans has encountered difficulties in replicating key findings from animal studies, such as the relationship between habit strength and the extent of prior training (de Wit et al. 2018; Gera et al. 2023; Pool et al. 2022). This difficulty is unsurprising, as humans are more likely to strategically adjust their behavior in response to shifting reward conditions (Seger and Spiering 2011). Nevertheless, a pioneering neuroimaging study that inducing habits in humans successfully replicated animal findings using overtraining procedures (Tricomi, Balleine, and O’Doherty 2009). This and other studies pinpoint the posterior putamen as a critical hub to learn and retrieve habitual behaviours (Tricomi et al., 2009; de Wit et al., 2012; McNamee et al., 2015). Trying to decipher the cortico-subcortical connectivities, de Wit et al. (2012) reported an increased anatomical pathway from premotor cortex to posterior putamen associated with individual’s tendency to habit-like performance. Yet, no information is available on the brain network involved in learning and overriding habitual actions. Deciphering whether similar or different neural circuits operate in expressing or reversing habits will offer a holistic view on how the habitual brain is organized (rather than isolated neural hubs) and provide insights to pathologies that implicate the habitual pathways.

Importantly, it remains unclear whether previous studies are truly showing the functioning of the habit system, especially when failing to demonstrate the expected overtraining effect. Recently, Luque et al. 2020 showed results in line with the expected overtraining effect. They manipulated the amount of training within subjects (overtrained vs. standard-trained stimuli) following a three-day training regime. Habit formation was assessed through a partial reversal test (in which one outcome was devalued) using time pressure (500ms to respond) which could take place either on the first day (standard-trained condition) or on the third day of the experiment (overtrained condition). For the habits test, the outcome of the habitual response was devalued (outcome devaluation test, Dickinson, 1985) so participants had to change their habitual response for still getting a valuable outcome. The protocol reveals additional response time (RT) costs when switching habitual responses during devaluation (RT Switch cost) (Luque et al. 2020). Importantly, this RT Switch cost was higher in overtrained conditions—condition in which habits are thought to be stronger producing more interference over new goal-directed responses. These findings have since been replicated by an independent pre-registered study (Nebe et al. 2024). The increased RT cost suggests greater difficulty in modifying established habitual responses and provides a novel measure to account for habit strength in humans. Luque et al.’s test reflects the common situation in which the two decision systems compete and the more advantageous goal-directed is finally expressed. Critically, this allows the study of the habit system without the need of artificially forcing habitual errors.

We aimed to expand our understanding of how habits are formed by modeling the learning process as a gradual accumulation of evidence (with drift diffusion modelling, DDM). In the context of habits, DDM can capture how repeated behaviors become faster, more consistent, and less dependent on conscious deliberation. In other words, habits shall rely on a higher drift rate, meaning that evidence accumulates more quickly toward a decision (Zhang et al. 2024). This reflects the efficiency of habits, as they rely on well-learned associations rather than slow, effortful reasoning. Another way habits shall manifest in a DDM is through a biased starting point. When a response is strongly habitual, the decision process may already be predisposed toward that option from the outset, making it more likely to be selected even if another response might be more optimal in a given situation. This bias reflects the automatic pull of habitual tendencies, which can be difficult to override. Overall, DDMs provide a useful framework for illustrating how habitual behaviors emerge, persist when no longer need and become automatic, ideally suited for the purpose of the present study.

We hypothesize that extensive training will increase RT Switch cost by changes in such drift diffusion computational processes. At the neural level, extensive training and habitual overriding produced in the outcome devaluation test are expected to recruit similar but divergent cortico-striatal habit circuitries (fMRI experiment). Moreover, where in the brain habits are stored will be leveraged with neuromodulation to provide a causal role of the prefrontal cortex in habit expression (transcranial magnetic stimulation (TMS) experiment).

## Results

### Habit Learning Dynamics

A reward learning task was employed (**Figure 1A**) with 33 participants in the fMRI experiment and 26 in the TMS experiment. Participants were trained over three days on stimulus-response links, with some stimuli receiving extensive training (overtrained) and some receiving standard training. On the final day, training and devaluation blocks were conducted inside the scanner. As expected, training led to improvements in establishing the correct associated answer to each cue (i.e. increased accuracy) and its speed when selecting the action (i.e. faster reaction times, RT). Consistent with optimal learning, accuracy during training was higher for overtrained stimuli compared to standard-trained stimuli (Estimate = 0.077, SE = 0.010, z = 7.853, p < .001; **Figure 1B**). Devalued trials revealed a reduction in accuracy for overtrained stimuli (i.e., fewer correct response switches) compared to the training phase, as indicated by a significant interaction between block × amount of training (F(1, 4850.35) = 22.981, p < .001; **Figure 1C**). These findings support our hypothesis that modifying habitual responses is more difficult in the overtrained condition.

**Figure 1.**
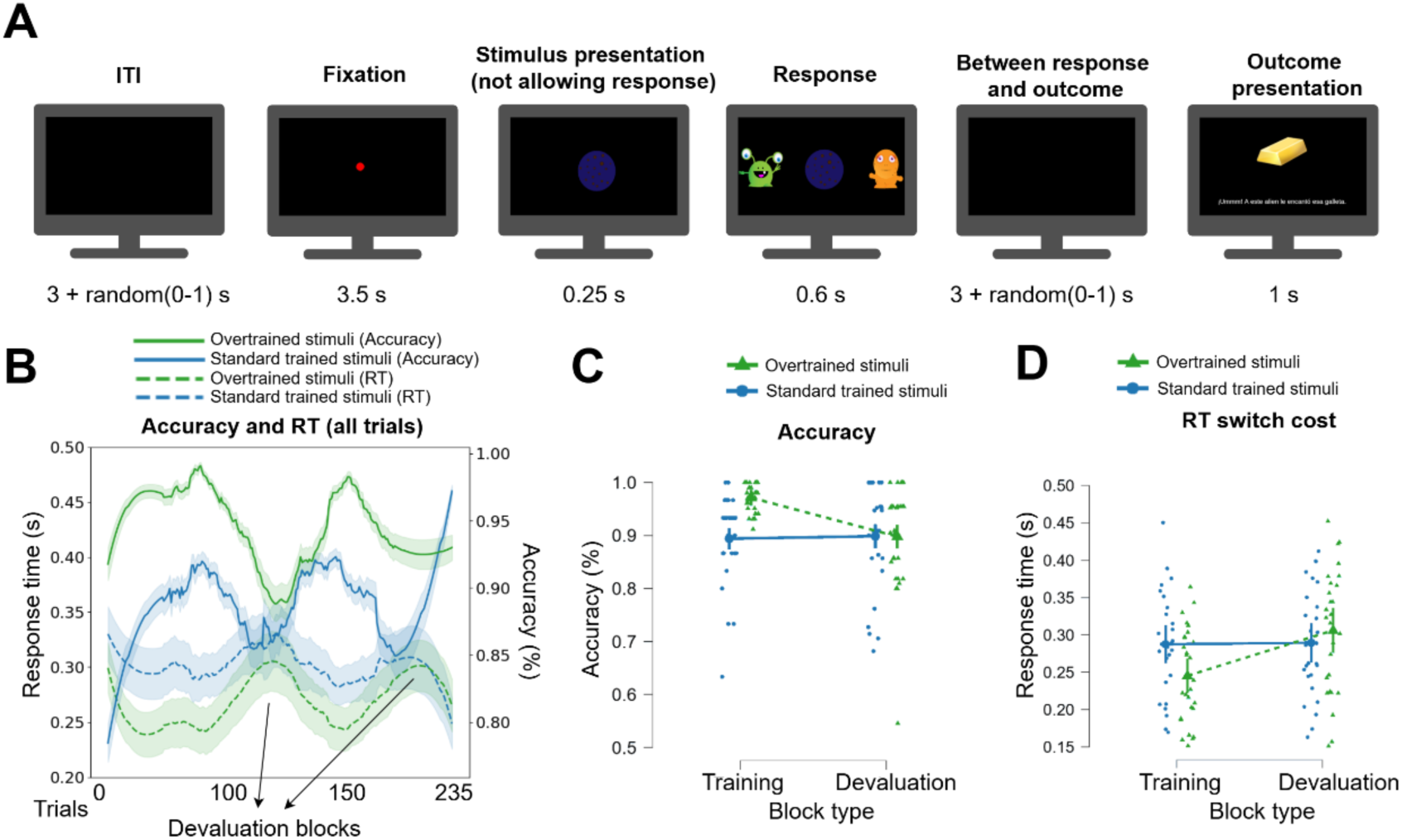
The Reward Learning Task and Behavioral Results. **A)** Example of a single trial. **B)** Accuracy and response time across trials, with a Savgol filter (window size = 71) applied to smooth the data for better visualization. Deviation width was reduced to 5% for clarity; shaded areas represent standard deviation. **C)** Comparison of performance between overtrained and standard-trained stimuli in terms of accuracy and **D)** RT switch cost during training and devaluation blocks.

Further analysis of learning dynamics revealed that response speed during training was faster for overtrained stimuli compared to standard-trained stimuli (Estimate = −0.045, SE = 0.004, z = −10.103, p < .001; **Figure 1D**). On devaluated trials, participants showed longer RTs for response switches in the overtrained condition compared to the training phase (interaction block × amount of training: (F(1, 27.04) = 20.936, p < .001; **Figure 1D**). These findings suggest that modifying a habitual response incurs greater cognitive costs in terms of RT for overtrained stimuli. These results are consistent with our hypothesis and previous findings demonstrating that RT Switch costs are greater for extensively trained stimuli.

### The Role of the Sensorimotor Circuit in Habit Expression and Goal-Directed Response Switching

Bayesian statistics with multiple comparison correction was used in both univariate and g-PPI connectivity analyses. Results reveal in the training phase for overtrained stimuli—compared to standard-trained stimuli—activity in the primary and secondary somatosensory cortex (bilateral), right superior temporal gyrus, and left primary motor area (**Figure 2A, Table S1A**). Whole-brain analysis did not reveal habit-related striatal activation. However, given the *a priori* significance of this area and following previous research (see Guida et al., 2022), we implemented a small volume correction (SVC) analysis using a striatum region of interest (ROI), which revealed bilateral activation in the posterior putamen (**Figure 2A, Table S1A**). To further investigate neural activity associated with goal-directed processes during training, we analyzed the inverse contrast (standard > overtrained stimuli). This analysis revealed activation in the bilateral posterior caudate, bilateral insula, left inferior parietal lobule, left dorsolateral prefrontal cortex (dlPFC), left orbitofrontal cortex (OFC), and left fusiform area (**Figure S1A; Table S1B**).

**Figure 2.**
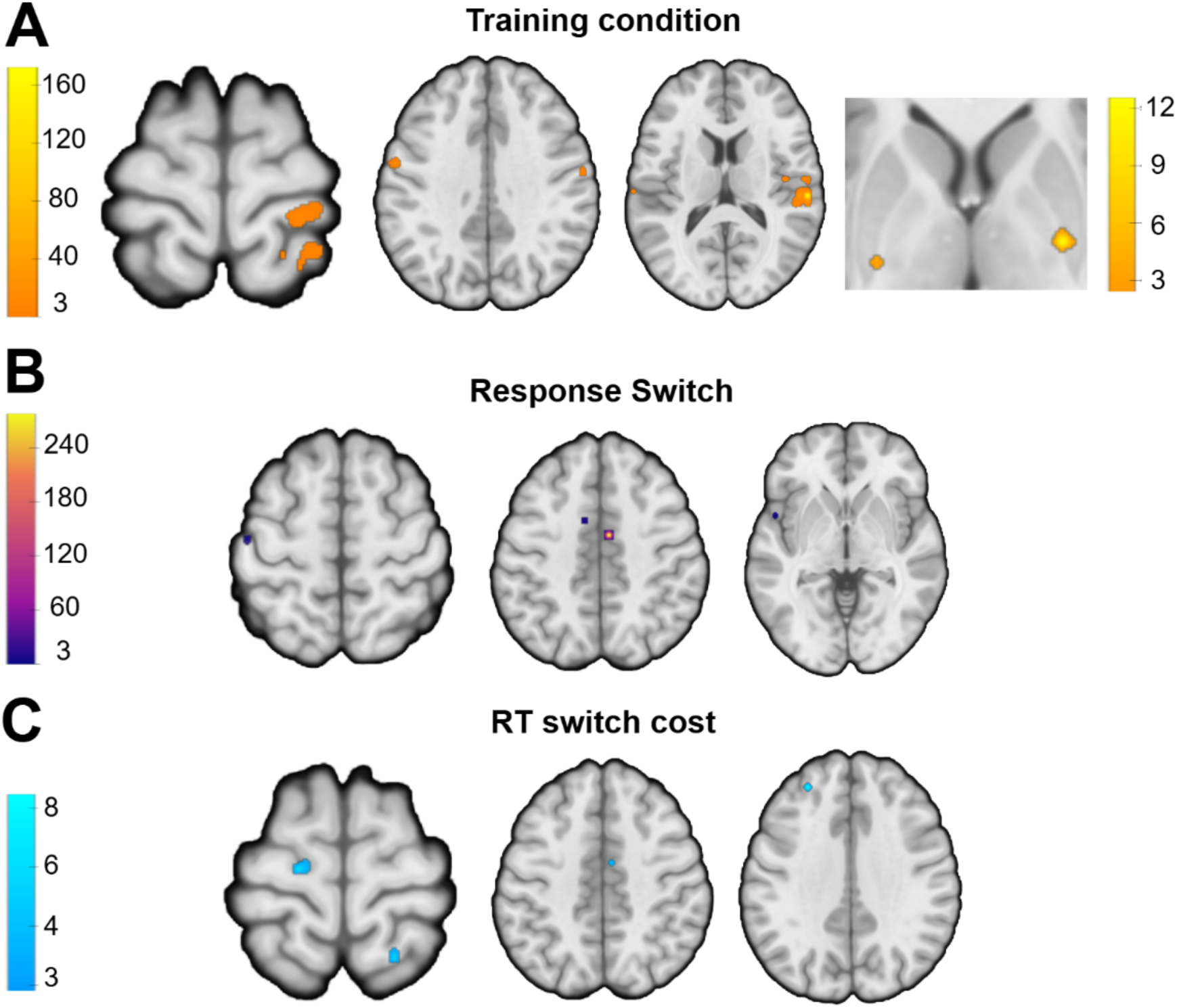
Brain Activity During Training and Devaluation in Habitual Behavior (Overtrained vs Standard Training). A Bayes factor >= 3 was used for thresholding. **A)** GLM univariate contrasts showing activation of habitual brain areas during training. Striatal activation in the right-most axial view was identified using a small volume corrected analysis with a striatal ROI (color scale shown on the right). **B)** GLM univariate contrasts highlighting activation of habitual brain areas when switching the overtrained S-R associations, and **C)** incorporating RT switch costs as a regressor to model BOLD data. For the complete pattern of results, see supplementary material Tables S1 to S3. No cluster extent threshold was applied, except for the first contrast, where single-voxel clusters were removed for visualization purposes due to the higher number of trials.

During the devaluation phase, neural activity related to the intrusion of well-learned habits (Response switch overtrained > standard trained) revealed activation of the left secondary motor cortex, left superior parietal lobule, premotor cortex, and motor cortex (**Figure 2B, Table S2A**), suggesting habitual activity from the sensorimotor circuit. For the inverse contrast (Response switch standard trained > overtrained), we observed activation in the bilateral caudate, right vlPFC, and right hippocampus/amygdala (**Figure S1B, Table S2B**). To further examine the neural correlates of RT switch costs as habits wore off, we regressed the RT costs (RT switch cost overtrained > standard trained), which revealed activation in the left PMC, right superior parietal, left dlPFC, and mid-cingulate (**Figure 2C, Table S3A**). No striatal activation was observed in either whole-brain or ROI analyses. The inverse contrast (RT switch cost: standard trained > overtrained) showed activation in right V1, left dlPFC, right ventrolateral prefrontal cortex (vlPFC), left inferior temporal cortex, right fusiform area, and right hippocampus/amygdala (**Figure S1C, Table S3B**). ROI analysis of the striatum additionally revealed activation in the bilateral caudate, left pallidum, and right anterior putamen (**Figure S1C, Table S3B**). At feedback, expected findings were obtained during reward receipts including anterior cingulate and amygdala for overtrained stimuli (**Table S8**).

### Habit Hubs Activity Explains the Degree of Switch Costs

Individual differences in habit expression and modulation have been shown along cortico-subcortical hubs. Individuals more prone towards goal-directed behavior largely depend on fronto-parietal regions (unlike those showing greater reliance on habitual behaviours) (Gera et al., 2023), while stronger tracts between left premotor and posterior putamen predicted greater habit perseverance (de Witt et al., 2012). As individuals may engage in different strategic forms, we hypothesized that brain activity in areas engaged with habitual responses (during S-R training) would be associated with the individual magnitude of RT Switch costs (at devaluation). To test this, we correlated specific brain activity from the habitual contrast in the training phase (overtrained > standard trained) with RT switch cost (baselined with respect to training performance). This analysis was conducted across all regions identified in the univariate contrasts to explore how training-related performance may impact subsequent habitual behavior in the devaluation phase. No significant correlations survived multiple comparison corrections (Bonferroni, False Discovery Rate) likely due to the stringent corrected significance threshold required for testing multiple ROIs (>25). Consequently, some significant correlations may not have been detected. Therefore, we also report findings based on uncorrected p-values. We observed that participants with greater activation in the left posterior putamen exhibited more habitual behavior in the devaluation phase (i.e. larger RT switch cost) (r = 0.48 p = 0.01; **Figure 3A**), suggesting that higher left putaminal activation may be associated with greater intrusion of habits. Interestingly, the opposite pattern was observed for standard-trained stimuli (**Figure 3A**; r = −0.36; p = 0.06). A similar pattern was found in the right secondary somatosensory cortex, but only for standard-trained stimuli (r = 0.44; p = 0.02; **Figure 3A**). This may suggest that higher somatosensory cortex activity during training facilitates the better encoding of less trained (generally weaker) associations, leading to a higher switching cost during devaluation. Conversely, in the left OFC, participants with higher activation during training for overtrained stimuli demonstrated greater flexibility and less habitual behavior during devaluation (r = −0.38; p = 0.04; **Figure 3A**), suggesting a stronger engagement of goal-directed processes in the OFC.

**Figure 3.**
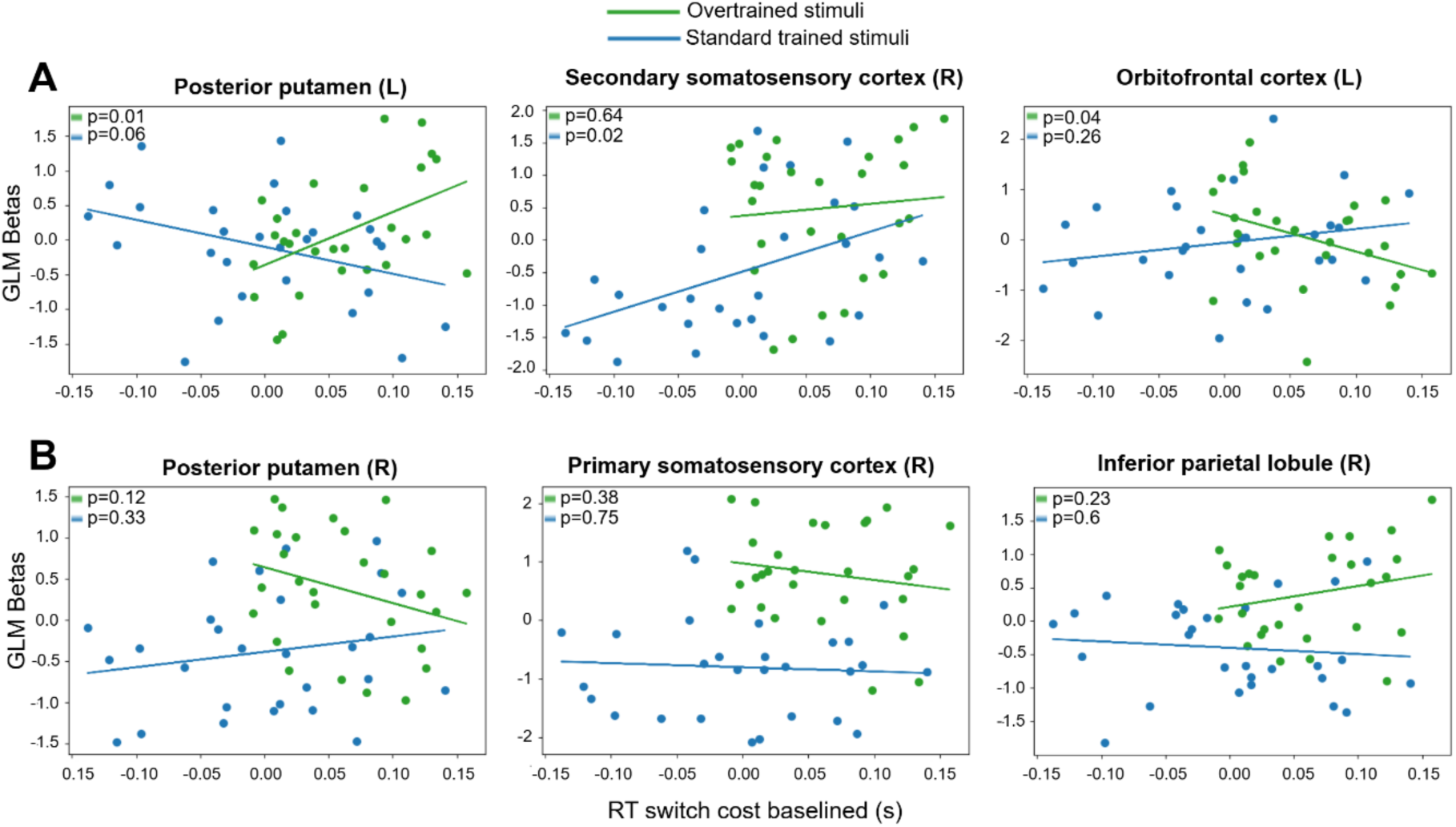
Brain activity during training correlates with switching during devaluation. Brain beta values during training (overtrained> standard trained) correlated with RT switch cost (uncorrected). **A)** Brain regions showing significant and **B)** non-significant correlations with RT switch costs, displaying the p-values for both overtrained and standard-trained conditions.

### Cortico-Subcortical Interactions when Expressing and Modulating Habits

To date, no clear habitual circuitry is available in the literature nor how transitions to new networks occur when overriding habits is required. To investigate how functional connectivity shifts during habitual behaviors and their modulation (i.e., goal-directed switching), we selected different seed regions for overtrained against standard-trained trials (using g-PPI): the right primary somatosensory cortex (S1) for the training condition, the motor cortex and left orbitofrontal cortex (OFC) for the switching condition, and the left premotor cortex (PMC) and right secondary somatosensory/inferior parietal lobule (S2/IPL) for the RT switch cost regressor. During training, using the right S1 as the seed region revealed significant connectivity with the left anterior insula, left OFC, and left DLPFC (**Figure 4A, Table S4A**). When switching overtrained responses (compared to switching standard trained responses), the motor cortex seed showed a functional connection with the right hippocampus/amygdala (**Table S4B**). This contrast also revealed connectivity between left OFC (seed) and right vlPFC (**Figure 4B, Table S5**). In the RT switch cost analysis, we observed significant connectivity between the left PMC and left putamen, as well as between the right S2/IPL and left SMA (**Figure 4C, Table S6**). These interactions are summarized in **Figure 4D** for clarity.

**Figure 4.**
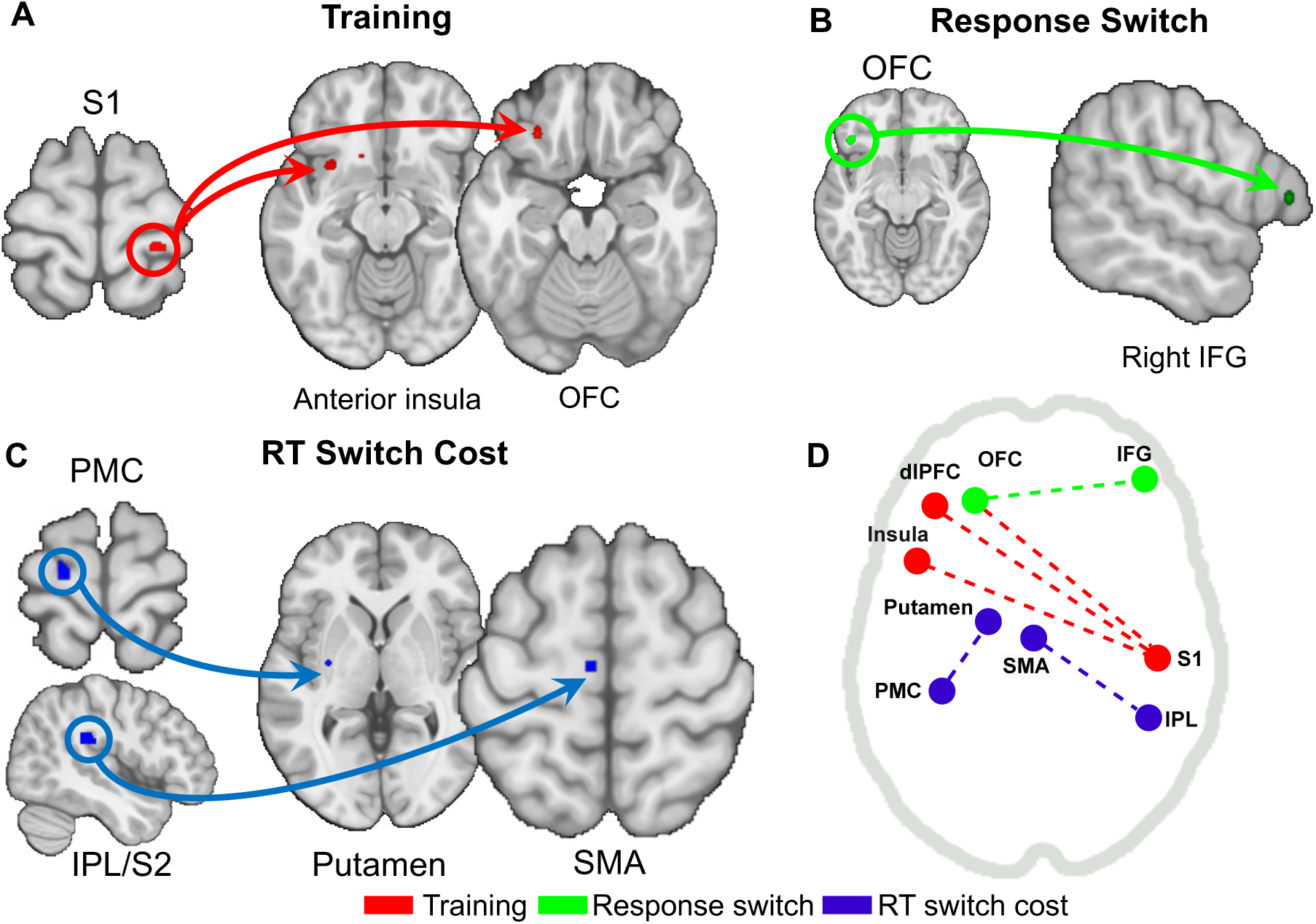
Functional connectivity findings in habit expression and modulation. Brain-wide connectivity patterns associated with **A)** training**, B)** response switching, and **C)** RT-switch cost conditions. The diagram illustrates interactions between regions identified in the univariate contrasts (training, switching, and RT Switch cost) as revealed by the g-PPI analysis. **D)** Summary of the main connectivity findings across conditions for each specific contrast with significant connectivity.

### Drift diffusion Processes in Habit Expression and Modulation

The final session of the fMRI experiment was used to fit using HDDM to both training and devaluation blocks. Following the model fitting, a posterior predictive check confirmed that the complete model —including all variables—successfully replicated the behavioral data across all conditions (training and devaluation blocks for both standard and overtrained stimuli; **Figure S2**). To assess model performance, we fitted five alternative model variations, each with a reduced set of parameters (separating conditions for *t*, *a*, *v*, and *t* = 0). Model comparison based on the DIC indicated that the complete model (*DIC* = −9054.27), which accounted for training and devaluation blocks as well as overtrained and standard-trained stimuli across all parameters (*t*, *v*, *a*), provided the best fit (**Table S7**). However, parameter recovery analysis revealed non-singular parameters, indicating that multiple parameter combinations could produce equivalent model fits in terms of global likelihood. While this limits direct interpretation of individual parameters, the successful fit to a drift diffusion model suggests that our study captured systematic changes in decision-making processes, consistent with an accumulation-to-threshold mechanism typical of automated behaviors (Zhang et al. 2024).

### Causal Contribution of the Premotor Cortex to Habits

To establish a direct link between our imaging findings and the behavioral activation of habits, we selectively modulated the left PMC (**Figure 5A**) to induce changes in either habit expression (training) or modulation (devaluation). The PMC is a key cortical area engaged in habitual control connected to the putamen (de Witt et al., 2012) and a putative target to modulate the expression of habits. The overall results (i.e., without distinguishing between real and sham conditions) replicated the overtraining effects observed in the fMRI experiment in terms of accuracy and RT Switch cost (see Supplementary Material). Importantly, we observed a global improvement in accuracy when real TMS was applied (compared to sham) as indicated by a significant main effect of TMS (*F*(1, 11,718) = 25.781, *p* < .001; **Figure 5B**, **5C)**. This enhancement was present across task conditions, suggesting that training was performed at a higher level following real TMS compared to sham stimulation **(Figure 5B**), an effect that also extended to devaluation tests (**Figure 5C**). This dual effect has important implications on how the brain executes habits, indicating that the left PMC role in habits is extensible to situations when habits are successfully overridden by the goal-directed system. That is, habitual control applies in the expression and the modulation of a habit, a function where the left PMC plays an essential biological function in both conditions. However, this improved accuracy did not result in significant differences in reaction times during either training (**Figure 5D**) or devaluation (**Figure 5E**). To control for potential motor side-effects of stimulation, a finger-tapping task was conducted, revealing a significant main effect of hand (faster responses with the right hand; *p* < .001) but no significant interaction between TMS and hand dominance.

**Figure 5.**
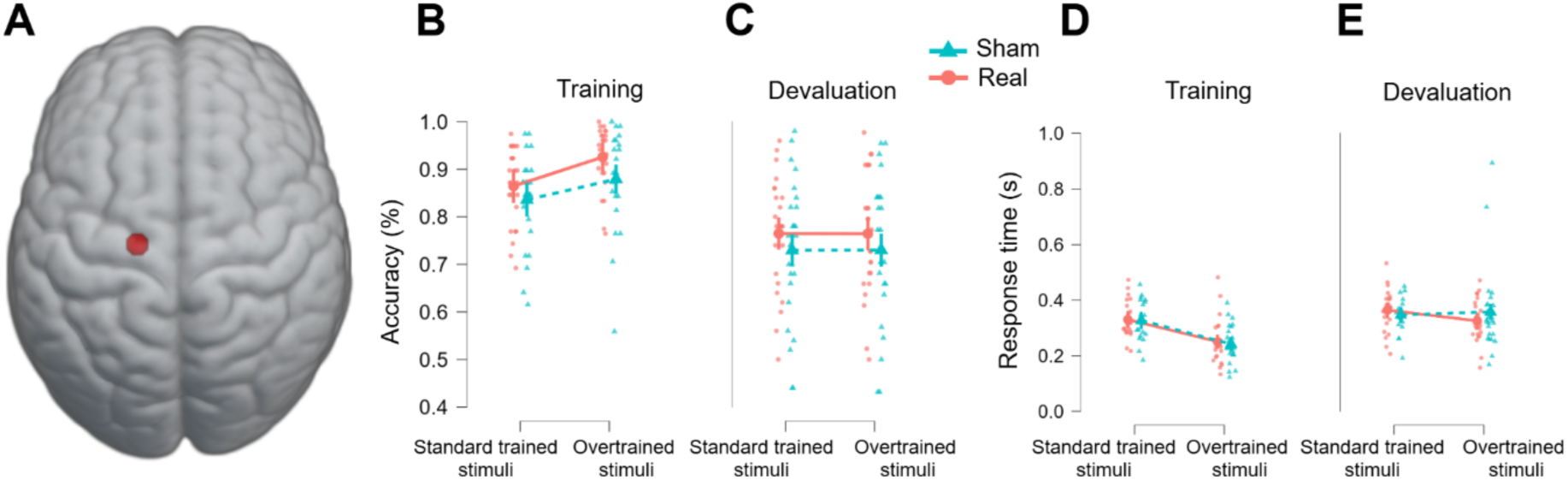
Behavioral findings and Neuromodulation (Real vs. sham TMS) for Overtrained and Standard-trained Stimuli. **A)** TMS stimulation target identified from GLM analysis, highlighting the left PMC; **B)** Behavioral results following TMS, showing accuracy in real vs sham conditions in the training; and **C)** devaluation conditions. **D)** RT in training and **E)** during the devaluation phase for both stimulus types.

## Discussion

Recent studies have failed to consistently replicate the expected relationship between the amount of training and habit strength, limiting translational research between human and animal models. In this study, we addressed these issues by assessing habit strength based on its interference with ongoing, incompatible goal-directed actions—specifically when participants were required to change their habitual responses under time pressure. The effectiveness of our paradigm stems from the focus on measuring response conflicts rather than direct choices (or the absence of choice), as is typically done in conventional devaluation methods (de Wit et al. 2012). The switch cost measure (with response-time constraints) effectively captured increased habit strength by revealing delayed response times when participants were required to break established S-R associations (Luque et al. 2020; Nebe et al., 2024). Additionally, the HDDM model decomposition of our behavioral data aligns with the notion that training leads to rapid response selection (Ashby, Turner, and Horvitz 2010; Daw, Niv, and Dayan 2005; Zhang et al., 2024). These findings suggest that drifts and evidence accumulation processes become automatized during habit expression and modulation (Ratcliff & Frank, 2012). Our results confirm that training, coupled with time pressure, plays a critical role in limiting goal-directed control, enabling a clearer manifestation of habitual tendencies (Hardwick et al., 2019).

The mechanisms underlying how the human brain navigates habits and shifts from goal-directed to habitual behavior remain unclear. Building on current behavioral theories concerning the transition from habitual to goal-directed control, our findings suggest the potential joint processing of habitual and goal-directed cues. In line with several studies, we observed activation in the sensorimotor cortex and putamen during training (Tricomi, Balleine, and O’Doherty 2009; de Wit et al. 2012; Delorme et al. 2016; Gera et al. 2023). However, this pattern is not always reported in the literature (see Valentin, Dickinson, and O’Doherty 2007; Watson, van Wingen, and de Wit 2018). We propose that the robust evidence for the role of the putamen in activating habitual behavior in our study stems from the strong habitual component of actions, which was promoted by both the training regime and the imposed response time constraints. The engagement of the cortico-subcortical neural system was further reflected in the strong effect of overtrained stimuli during training, which importantly induced longer switching costs (demonstrated by positive correlations with both putamen and S1). This result confirms a critical link between sensorimotor control during training and its subsequent impact when habits must be overridden, highlighting a dynamic interaction between habitual and goal-directed processes.

When behavior operates in habitual mode, control-related brain areas likely monitor signals of updated contextual information to determine whether to maintain or override ongoing habits. The best-fitting DDM model showed differences in diffusion and threshold variables in a similar manner for both training and devaluation blocks, further supporting this notion. Moreover, the primary somatosensory cortex (S1) contributes to habit organization by interacting with regions typically associated with goal-directed control, such as the anterior-insular cortex (Osada et al. 2024) and visual cortex (Cruz et al. 2023). Similarly, the observed connectivity between the secondary somatosensory cortex (S2)/inferior parietal lobe (IPL) and the SMA, as well as between the M1 and the hippocampus/amygdala, underscores the complex interplay between motor execution, planning, and memory processing when behavior is governed by habitual control. These findings suggest that the somatosensory cortex is part of a broader network involved in integrating sensory inputs and emotional salience with ongoing motor responses. This network is crucial for both habitual and goal-directed control, facilitating the coordination of actions based on past experiences and current goals (Pantoja et al. 2007; Pleger et al. 2009).

A longstanding question in the literature is how the brain overrides habitual behaviors. Identifying the neural mechanisms that disengage habits when they are no longer needed is critical for understanding the flexibility of the habitual system and when this becomes dysfunctional (as in pathological conditions such as addiction or OCD). Our findings provide evidence for a predominantly motor-related cortical network involved in both the expression and modification of overtrained habits, including the left PMC, S2/IPL, left SMA, IFG, and cingulate cortices. In particular, connectivity analyses highlighted the interaction between the PMC and putamen, supporting previous findings linking putaminal activity to habitual behavior (de Wit et al. 2012). However, given that habits are no longer adaptative during devaluation (i.e., when they begin to wear off), both goal-directed and habitual control mechanisms require co-existence in the brain to successfully guide adaptive behaviour. We have identified a key transition from habitual behavior, mediated by the right IFG — previously implicated in behavioral control (Cieslik et al. 2013) — and the middle cingulate cortex, which is involved in motor planning and adaptive control (Hoffstaedter et al. 2014). The IFG, in particular, is strongly associated with goal-directed processes, suggesting its involvement in both habitual and goal-directed actions (or the transition between the two). These regions appear to be essential for overriding established habitual responses and represent potential targets for neuromodulation protocols in disorders characterized by the overexpression of habits.

Regarding the cingulate cortex, a large body of literature has linked this region with habitual behavior, with studies implicating different sections along the anterior-to-posterior axis depending on the specific function examined (Liljeholm, Molloy, and O’Doherty 2012; Liljeholm, Dunne, and O’Doherty 2015; Eryilmaz et al. 2017; Watson, van Wingen, and de Wit 2018). In our study, the posterior cingulate was engaged during training in the overtrained condition, whereas habit switching elicited a more anterior activation pattern (Watson, van Wingen, and de Wit 2018). This suggests that cingulate involvement in habit expression and modulation may occur at distinct anatomical locations, depending on the behavioral context. Critically, no striatal activation was observed in the overtrained conditions during devaluation, either in whole-brain or ROI analyses. One intriguing possibility is that extended training facilitated a transfer of control from the basal ganglia to cortico-cortical projections, consistent with the neurobiological model of automaticity proposed by Ashby et al. (2007). Under this framework, as habits become ingrained, their expression may rely more on cortical connectivity, particularly in novel situations where habitual responses must be overridden. However, despite this shift, residual subcortical activity may continue to exert influence, leading to intrusions from previously reinforced habit-related regions.

To investigate the cortical role in habit expression and modulation, we applied cTBS (inhibitory) over the left PMC (comparing its effects to a sham condition). Overall, cTBS led to an improvement in accuracy across both training and devaluation blocks. This improvement was likely driven by the inhibition of habitual circuitry, prompting participants to adopt a more goal-directed and controlled response strategy—even during training, a phase typically observed to foster habitual control. At the physiological level, the presumed reduction in PMC excitability might have increased the influence of goal-directed decision-making processes, thereby improving accuracy. This interpretation is consistent with the role of the posterior putamen-PMC pathway in habit formation (de Witt et al., 2012) and with previous accounts suggesting how PMC stimulation might modulate putaminal activity (van Holstein et al. 2018), effectively suppressing habit-related signals associated with well-learned cues. Compatible with the above interpretation, the local effects of PMC stimulation may have enhanced participants’ overall cognitive control or attentional mechanisms (similar to Rowe et al. (2002). This interpretation aligns well with research suggesting that the PMC is involved in cognitive control processes, including the selection and initiation of actions based on current goals (Gremel and Costa 2013). In contrast, an alternative explanation — that PMC stimulation enhances putaminal functions — is less probable, as habitual behaviors typically plateau with consistent training, limiting further improvements in performance. Overall, both options are plausible, given that a reduction in putamen activity during habitual behavior may have disrupted the balance between habitual and goal-directed processes, potentially increasing cognitive control demands. Altogether, the PMC seems to be involved in both the expression and overriding of habits, thereby suggesting its broader role when management of habit cues are critical to adapt behaviour.

## Methods

### Participants

Experiment 1 (fMRI study) initially recruited 33 participants from the Universidad Autónoma de Madrid (mean age = 20.6 years, SD = 3.2; 29 right-handed; 22 female). As an inclusion criterion, participants were required to have no history of neurological disorders, as confirmed through self-reported questionnaires. Individuals with mild-to-severe neurological conditions were excluded from the study, including those with moderate disorders requiring medication. Four participants were excluded from the final analysis: three due to excessive head movement during the fMRI session (see fMRI analysis section for details) and one due to low performance in the devaluation block. The final analyzed sample consisted of 28 participants.

For Experiment 2, 26 participants were recruited from the Universidad Complutense de Madrid (mean age = 23.0 years, SD 4.5; all right-handed; 20 female) to complete two TMS sessions (real and sham). To ensure that participants fully understood the devaluation procedure, each block included three validation trials in which they freely chose between the rewards. Given that Experiment 2 involved four devaluation blocks (due to the inclusion of both real and sham TMS sessions), compared to only two in Experiment 1, we adjusted the exclusion criteria. Participants were excluded only if they committed ≥1 consumption trial error in ≥2 devaluation blocks (instead of ≥1 error in a single block, as in Experiment 1). Based on this criterion, one participant was excluded, resulting in a final sample of 25 participants. All participants provided written informed consent in accordance with the Declaration of Helsinki. The institutional Ethical Committee of HM hospitals approved the experimental protocols.

#### Reward Learning Task

A reward learning task (Luque et al. 2020) was used across studies. Each trial began with a fixation cross (500ms), followed by the presentation of a stimulus (an image of a cookie) at the center of the screen, with two alien characters positioned on each side (Figure 1). Participants were instructed to press the <q> key to give the cookie to the left alien or the <p> key to give it to the right alien. Each alien preferred only one type of cookie; selecting the correct option resulted in a favorable outcome (gold or diamond), while an incorrect choice led to an unfavorable outcome (coins). Participants were required to respond within a 600 ms time limit; failure to do so resulted in a timeout screen, invalidating the trial. This time constraint was designed to engage the habitual system by limiting the time available for goal-directed decision-making (Keramati, Dezfouli, and Piray 2011). The task included four different types of stimuli (cookies), distinguished only by color (brown, blue, purple, and yellow). The alien images were included purely for representational purposes, serving as a cover story to facilitate memory for the associations rather than requiring participants to make arbitrary left or right choices.

During the outcome devaluation tests, one of the two positive outcomes (gold or diamond) was devalued in each devaluation block, with the specific devalued outcome counterbalanced across participants. All other rewards retained their original value. As a result, participants had to override their habitual response to avoid selecting the now-devalued outcome. Note that the devaluated outcome could be obtained through two different stimulus-response associations: one involving the overtrained stimulus and another involving the standard-trained stimulus. This arrangement allowed for a direct comparison of the effects of training on behavior during the devaluation phase. Additionally, it enabled a comparison of devalued versus still-valued trials. Experiment 2 followed the same devaluation block structure per session; however, participants completed two separate sessions (real and sham TMS), conducted at least one week apart.

#### Experimental Design

The reward learning task was employed with specific adaptations to meet the requirements of each study while maintaining identical outcome measures. Experiment 1 consisted of three consecutive sessions conducted over three days. The first two sessions were completed online, while the final session took place entirely inside the scanner, including both training and devaluation conditions. After the first session, participants’ accuracy was assessed, and those scoring ≤70% were excluded from further participation. Each online session included four training blocks, with a total of 44 trials per block: 34 trials featuring overtrained stimuli and 5 trials featuring standard-trained stimuli, amounting to 396 trials in total. Additionally, a devaluation block of four trials was introduced between the third and fourth training blocks. The inclusion of the devaluation block served to ensure that participants paid attention to the stimulus-response (S-R) associations that yielded the best rewards. This design reinforced the need to recall specific S-R pairings rather than relying solely on simple left-right decision-making strategies.

The final session was conducted inside the scanner. Participants first completed two training blocks with the same configurations as in the previous sessions. Next, a devaluation block with 44 trials (11 trials per condition: overtrained and standard-trained stimuli) was introduced, in which one of the trained associations (overtrained or standard-trained) was devalued (counterbalanced across participants). To avoid confusion between devaluation effects and regular outcomes, another training block followed, using the same configurations as the previous training blocks. Finally, a second devaluation block was introduced, devaluing the other condition (overtrained or standard-trained), while the previously devalued stimulus maintained its original value.

Experiment 2 consisted of two neuromodulation sessions (real and sham TMS) conducted in the laboratory using a within-subject design. Before each laboratory TMS session, participants completed two online sessions, followed by a final session in the laboratory. These sessions were conducted on consecutive days for each TMS condition. The task conditions remained identical to those described in the previous section.

#### Behavioral Analysis

A linear mixed-effects model (LMM) was used to test the statistical significance of the behavioral effects across both experiments.

For the response time (RT) analysis, we implemented a maximal random effects structure justified by our experimental design, as follows:

*response_time ∼ block_type * overtraining_stim + (1 + block_type * overtraining_stim | participant)*

From now on, we will refer to this model as the RT switch cost paradigm.

For the response accuracy analysis, the model was structured as follows:

*response_accuracy ∼ block_type * overtraining_stim + (1|participant)*

In Experiment 2, RT was analyzed as follows:

*response_time ∼ block_type * is_overtraining_stim * condition + (1|participant).*

The model for accuracy was defined as:

*response_accuracy ∼ block_type * is_overtraining_stim * condition + (1|participant)*

#### fMRI Univariate Contrasts

Univariate contrasts were computed using nilearn 0.4.2 (Abraham et al. 2014) from stimulus onset (zero duration). Multiple univariate contrasts were conducted to account for the different task conditions:

##### a) Training phase

During the training phase, we were interested in observing the learning differences between the overtrained and standard-trained stimuli. To achieve this, we compared responses when overtrained stimuli were presented versus when standard-trained stimuli were shown (only considering the correct responses). This contrast is referred to as “*Response overtrained correct > Response not overtrained correct*.” We assume this contrast primarily reflects habitual processes, while goal-directed processes may emerge when the contrast is inverted.

##### b) Devaluation phase

The following contrasts were designed to assess the RT switch cost hypothesis, which has not been previously studied using fMRI. The initial contrast derived from this condition is *“Response switch overtrained > Response switch not overtrained.”* Note, however, that this contrast is not the most precise for testing the RT switch cost hypothesis since it includes all response switch trials, even those that did not show an additional RT cost. Nonetheless, due to the limited number of trials, this contrast provides a more reliable method for testing the hypothesis.

We also analyzed a more refined version of this hypothesis: *“RT switch cost baselined positive overtrained > RT switch cost baselined positive not overtrained.”* This contrast, a subset of the previous one, incorporates an RT baseline from the final training block (i.e., RTs from the last training block were subtracted from RTs in the devaluation blocks). Although this contrast includes fewer trials, applying Bayesian analysis allowed for reliable results that complemented the broader contrast. We assume both contrasts primarily reveal habitual activation, while goal-directed processes are expected when the contrast is inverted.

Bayesian statistics with multiple comparison correction was employed using the BayesFactorFMRI package in Python (Han 2021a) for both univariate and g-ppi analyses. The primary advantage of using Bayesian statistics is the enhanced sensitivity for studies with limited trials, such as our devaluation blocks (Han 2021b; Han and Park 2018; Magerkurth et al. 2015). Full technical details of the fMRI univariate analyses can be found in the Supplementary Material.

#### fMRI Connectivity Analysis: g-PPI

To examine cortico-subcortical interactions during habit expression, we conducted functional connectivity analysis using a generalized psycho-physiological interactions model (g-PPI) (McLaren et al. 2012). This analysis aimed to provide additional hints concerning whether a particular region is involved in habitual or goal-directed circuitry or whether there is an interaction between the two. Hence, the contrasts used for this analysis were the same as those described in the univariate contrasts section above. A detailed explanation of how the regressors were included can be found in the Supplementary Material.

#### Hierarchical Drift Diffusion Model (HDDM)

To investigate variations in response speed during the creation and expression of habits, we fitted a hierarchical drift diffusion model (HDDM) to account for all data from the training and devaluation blocks (Experiment 1). We chose a drift diffusion model (Ratcliff 1978; Ratcliff and Rouder 1998) to incorporate both response time and accuracy rather than focusing solely on accuracy, as in most general reinforcement learning algorithms. The model assumes that, over time, the evidence for choosing an answer accumulates, with the most prioritized option being selected (similar to theoretical accounts of habits; Ratcliff and Frank 2012). The rate of this accumulation process is characterized by the drift rate: the steeper the slope, the faster the decision. This feature is particularly useful for modeling habitual behavior (Ratcliff and Frank 2012). Further technical details can be found in the Supplementary Material.

#### TMS protocol

In Experiment 2, we aimed to target a key hub of the habitual circuitry using continuous theta-burst stimulation (cTBS) over the left PMC (technical details can be found in the Supplementary Material). The experimental procedure was divided into two phases. In the first phase, participants completed a training session at home over two consecutive days. On the third day, TMS was administered in either active or sham conditions (using a within-subject counterbalanced, single-blind design). Immediately after cTBS, participants engaged in a 5-minute finger-tapping task, which served as a control condition to ensure minimal interference with M1. Subsequently, they completed the reward learning task, including training and devaluation conditions. An interval of two to three weeks was introduced between the two phases to reduce the retention of associations formed during the first phase. This interval is longer than that outlined in the standard safety guidelines for TMS (i.e., a one-week interval between sessions) (Rossi et al. 2009).

During the second phase (real or sham, depending on the counterbalanced order), participants were exposed to novel stimuli characterized by different shapes and colors to ensure the formation of new associations. The TMS condition assigned to each participant in the second phase was the inverse of what they had received in the first phase, ensuring a counterbalanced design.

## Acknowledgments

We are grateful to Pasqualina Guida and David Mata-Marín (CINAC, Hospital Universitario HM Puerta del Sur, Móstoles) for their help in the fMRI data acquisition. MM was funded by Instituto de Salud Carlos III (PFIS contract) and Fundación HM Hospitales. IO was funded by Instituto de Salud Carlos III (Miguel Servet, CP18/00038). This research has been funded by ISCIII (PI19/00298) from the Ministry of Science and Innovation and PROYEXCEL_00287, awarded by the Junta de Andalucía (Spain). The funders played no role in the idea, design, data collection or analysis, decision to publish, or manuscript editing and writing.

## Author Contributions

M.M. design, data collection, analysis, writing; D.L. design, data collection, writing; I.O. design, data collection, writing.

## Competing Interests statement

The authors declare no competing interest.

## Data availability

Access to behavioural and fMRI data with scripts is available at https://osf.io/w86nt/files/osfstorage.

## Notes

https://osf.io/w86nt/

### Competing Interest Statement

The authors have declared no competing interest.

