## Supplemental material for "The Neural Basis of Habit Formation Measured in Goal-Directed Response Switching"

### Methods

#### *fMRI acquisition and preprocessing*

Imaging data was collected using a 3T hybrid PET-MRI scanner (mMR Biograph, Siemens AG, Germany) with a 12-channel head array coil. Task-based fMRI was acquired using a single-shot gradient-echo echo-planar imaging (EPI) 2D pulse sequence with the following parameters: TR/TE = 2000/30ms; optimum flip angle using the Ernst equation i.e., 79°, spatial resolution = 3x3x4mm3; field of view = 192mm; matrix = 64×64; slice thickness= 4mm and acceleration factor of 2 (IPAT2). Five fMRI runs were acquired per session (285 volumes per run, 12mins) with 1-min pause between runs. Imaging protocol also included a 3D T1-weighted MP-RAGE (TR/TE/TI =143 2300/3.34/900ms;flip angle = 8; and isotropic spatial resolution = 1mm3, FoV = 256mm, matrix = 256×256, slice thickness = 1mm); and a fieldmap generated from two 2D gradient-echo images (TR/TE1/TE2 = 455/4.92/7.38ms, flip angle = 60° with the spatial resolution as the fMRI EPI volumes). Results included in this manuscript come from preprocessing performed using fMRIPrep 20.2.1 (Esteban et al. 2019) (RRID:SCR 016216), which is based on Nipype 1.5.1 (Gorgolewski et al. 2011); RRID:SCR 002502).

#### Anatomical data preprocessing

A total of 1 T1-weighted (T1w) images were found within the input BIDS dataset. The T1-weighted (T1w) image was corrected for intensity non-uniformity (INU) with N4BiasFieldCorrection (Tustison et al. 2010), distributed with ANTs 2.3.3 (Avants et al. 2008), RRID:SCR 004757), and used as T1w-reference throughout the workflow. The T1w-reference was then skull-stripped with a Nipype implementation of the antsBrainExtraction.sh workflow (from ANTs), using OASIS30ANTs as target template. Brain tissue segmentation of cerebrospinal fluid (CSF), white-matter (WM) and gray-matter (GM) was performed on the brain-extracted T1w using fast (FSL 5.0.9, RRID:SCR 002823 (Zhang, Brady, and Smith 2001). Volume-based spatial normalization to one standard space (MNI152NLin2009cAsym) was performed through nonlinear registration with antsRegistration

(ANTs 2.3.3), using brain-extracted versions of both T1w reference and the T1w template. The following template was selected for spatial normalization: ICBM 152 Nonlinear Asymmetrical template version 2009c (Fonov et al. 2009), RRID:SCR 008796; TemplateFlow ID: MNI152NLin2009cAsym].

**Functional data preprocessing**

For each of the 5 BOLD runs found per subject (across all tasks and sessions), the following preprocessing was performed. First, a reference volume and its skull-stripped version were generated using a custom methodology of fMRIPrep. A B0-nonuniformity map (or fieldmap) was estimated based on a phase-difference map calculated with a dual-echo GRE (gradient-recall echo) sequence, processed with a custom workflow of SDCFlows inspired by the epidewarp.fsl script and further improvements in HCP Pipelines (Glasser et al. 2013). The fieldmap was then co-registered to the target EPI (echo-planar imaging) reference run and converted to a displacements field map (amenable to registration tools such as ANTs) with FSL's fugue and other SDCflows tools. Based on the estimated susceptibility distortion, a corrected EPI (echo-planar imaging) reference was calculated for a more accurate co-registration with the anatomical reference. The BOLD reference was then co-registered to the T1w reference using flirt (FSL 5.0.9, Jenkinson and Smith 2001) with the boundary-based registration (Greve and Fischl 2009) cost-function. Co-registration was configured with nine degrees of freedom to account for distortions remaining in the BOLD reference. Head-motion parameters with respect to the BOLD reference (transformation matrices, and six corresponding rotation and translation parameters) are estimated before any spatiotemporal filtering using mcflirt (FSL 5.0.9, Jenkinson et al. 2002). BOLD runs were slice-time corrected using 3dTshift from AFNI 20160207(Cox and Hyde 1997), RRID:SCR 005927). The BOLD time-series (including slice-timing correction when applied) were resampled onto their original, native space by applying a single, composite transform to correct for head-motion and susceptibility distortions. These resampled BOLD time-series will be referred to as preprocessed BOLD in original space, or just preprocessed BOLD. The BOLD time-series were resampled into standard space, generating a preprocessed BOLD run in MNI152NLin2009cAsym space. First, a reference volume and its skull-stripped version were generated using a custom methodology of fMRIPrep. Several confounding time-series were calculated based on the preprocessed BOLD: framewise displacement (FD), DVARS and three region-wise global signals. FD was computed using two formulations following Power (absolute sum of relative motions, (Power et al. 2014) and Jenkinson (relative root mean square displacement between affines, (Jenkinson et al. 2002)). FD and DVARS are calculated for each functional run, both using their implementations in Nipype (following the definitions by Power et al. 2014). The three global signals are extracted within the CSF, the WM, and the whole-brain masks. Additionally, a set of physiological regressors were extracted to allow for component-based noise correction (CompCor, (Behzadi et al. 2007). Principal components are estimated after high-pass filtering the preprocessed BOLD time-series (using a discrete cosine filter with 128s cut-off) for the two CompCor variants: temporal (tCompCor) and anatomical (aCompCor). tCompCor components are then calculated from the top 2% variable voxels within the brain mask. For aCompCor, three probabilistic masks (CSF, WM and combined CSF+WM) are generated in anatomical space. The implementation differs from that of Behzadi et al. in that instead of eroding the masks by 2 pixels on BOLD space, the aCompCor masks are subtracted a mask of pixels that likely contain a volume fraction of GM. This mask is obtained by thresholding the corresponding partial volume map at 0.05, and it ensures components are not extracted from voxels containing a minimal fraction of GM. Finally, these masks are resampled into BOLD space and binarized by thresholding at 0.99 (as in the original implementation). Components are also calculated separately within the WM and CSF masks. For each CompCor decomposition, the k components with the largest singular values are retained, such that the retained components' time series are sufficient to explain 50 percent of variance across the nuisance mask (CSF, WM, combined, or temporal). The remaining components are dropped from consideration. The head-motion estimates calculated in the correction step were also placed within the corresponding confounds file. The confound time series derived from head motion estimates and global signals were expanded with the inclusion of temporal derivatives and quadratic terms for each (Satterthwaite et al. 2013). Frames that exceeded a threshold of 0.5 mm FD or 1.5 standardised DVARS were annotated as motion outliers. All resamplings can be performed with a single interpolation step by composing all the pertinent transformations (i.e. head-motion transform matrices, susceptibility distortion correction when available, and co-registrations to anatomical and output spaces). Gridded (volumetric) resamplings were performed using antsApplyTransforms (ANTs), configured with Lanczos interpolation to minimize the smoothing effects of other kernels (Lanczos 1964). Non-gridded (surface) resamplings were performed using mri vol2surf (FreeSurfer). Several internal operations of fMRIPrep use Nilearn 0.6.2 ((Abraham et al. 2014), RRID:SCR 001362), mostly within the functional processing workflow. For more details of the pipeline, see the section corresponding to workflows in fMRIPrep's documentation.

Three participants were discarded for excessive head movements inside the scanner. The threshold for this was considered by using the framewise displacement (FD), which aggregates the 6 usual movement parameters (rotation and translation in their respective directions) obtained during the preprocessing steps. We discarded all the subjects that had an excessive movement in >= 2 blocks (of 5 in total), where we considered excessive movement when there were >= 30% of volumes with >= 0.2 mm FD. MRIQC (Esteban et al. 2017) was used in this step.

**fMRI analysis**

For univariate contrasts, we fitted a GLM including nine confounds columns extracted in the pre-processing steps (six for translation and rotation head movements, CSF, white matter, and global signal). Drift correction regressors were also included by applying a high pass filter at the standard cutoff frequency (0.01 Hz). An auto-regressive model (ar3) was used, taking the middle slice as the reference in time-space due to the re-alignment done in the pre-processing steps in fMRIprep. We applied the SPM HRF model, including the first derivative. Finally, a 5 $mm^3$ FWHM smoothing was applied at $1^{st}$ level analyses, and a 6 $mm^3$ FWHM for the $2^{nd}$ level. To increase the goodness of fit, for each contrast, we also included the regressors corresponding to other relevant conditions of the task (e.g., fixation, ITI, response, and feedback) with duration set to zero.

In addition to the whole-brain analysis, we also included ROI analysis. These were included due to the strong a priori hypotheses (Guida et al. 2022) to further explore the specific activity of the striatum. We used the Adult brain maximum probability map ("Hammersmith atlas"; n30r83) (Hammers et al. 2003) in MNI space to select the putamen section.

**fMRI connectivity analysis: g-PPI**

The g-PPI procedure is as follows: 1) a ROI seed is selected based on the significant results of the univariate analyses; 2) The de-meaned BOLD signal of the ROI (physiological regressor) is multiplied with the behavioral condition regressor (HRF convolved and de-meaned); 3) The psychological regressor for each condition of the contrast are included in a GLM. The parameters of the model are the same as in the univariate analyses described above; 4) Last, the GLM is computed in a whole-brain manner to check for significant interactions in any voxel (univariate-style).

**Computational model**

To fit the model, we separated the trials into four conditions: (i) training trials with overtrained stimuli and (ii) standard trained stimuli, (iii) devaluation trials with overtrained stimuli, and (iv) with standard trained stimuli. With this approach, the model provides separate estimates for each parameter depending on the condition, but still accounts for all the different conditions to maximize the data global likelihood when training the model (L(θ | Data, Condition). This implies that parameter estimates for one condition (e.g. training blocks) can influence the parameter estimates of the other conditions (e.g. devaluation blocks) and vice versa. The model takes into account all the data at once and finds the best parameters that explain the entire dataset, considering the block type. To this, we used the HDDM software in Python (Wiecki, Sofer, and Frank 2013), which allows flexible estimation of drift-diffusion models through hierarchical Bayesian parameter estimation. The model simultaneously estimates subject and group parameters, where individual subjects are assumed to be drawn from a group distribution (Pedersen, Frank, and Biele 2017).

**TMS protocol**

We employed continuous theta-burst stimulation (cTBS), which is well-known to reliably modulate motor cortical excitability for about 20-60 min post-stimulation (Huang et al. 2005). We used a 70 mm diameter figure-of-eight coil (Magstim Rapid2, Whitland, UK) to apply cTBS guided with a neuronavigation system (Brainsight; Rogue Research, Canada).

Before delivering neuromodulation, we first determined stimulation thresholds by delivering single pulses with the coil placed tangentially over the left M1, handle pointing 45° backwards and laterally. Motor evoked potentials (MEPs) from the left first dorsal interosseous were registered to detect the motor "hotspot", marked with the neuronavigation system. Then, the active motor threshold (AMT) was defined as the minimal output intensity producing MEPs of ~100 μV amplitude. To this, we used a threshold-tracking method with maximum-likelihood parameter estimation by sequential testing (PEST) software (MTAT 2.0). AMT was obtained with participants exerting minimal muscle activity by squeezing between forefinger and thumb below 100 μV (visually inspected).

Individual T1-weighted structural MRI scans were done on each participant to target the left PMC (MNI coordinates: [-18, -18, 68]; obtained from Experiment 1; **Figure 2C**), using Brainsight navigation system with T1s registered from MNI to each subject space. A Polaris infrared camera (NorthernDigital, Canada) tracked the markers on the subject's tracking glasses and the TMS coil in real-time (used also during the delivery of cTBS to reduce possible displacements). Coil orientation and positioning were marked with help from a neuronavigation device as to ensure minimal spatial variability. The co-registration between the participant's head and their MRI was achieved through identification of anatomical landmarks using Brainsight's pointer tool.

cTBS was delivered with 3 pulses at 50 Hz, repeated at 200 msec for 40 sec (600 pulses total) at 80% of participants' AMT over left PMC. Sham protocol was identical to real cTBS, but with the coil tilted 90° away from the scalp as to avoid any potential impact on participants' scalp. Maximum stimulator output was 39.2 ± 5.2 % for real cTBS and 39.9 ± 5.3 % for sham. Participants were screened for safety (Keel, Smith, and Wassermann 2001) and reported no side effects. Session order was counterbalanced across participants.

### Results

#### TMS behavioral results

As indicative of optimal learning (independent of stimulation), accuracy during training was higher for overtrained stimuli compared to standard trained stimuli (Estimate = 0.052, SE = 0.010, z =

5.428, $p < .001$; in both real and sham conditions). Further exploring the learning dynamics, we observed that RTs during training was faster for overtrained stimuli compared to undertrained trained stimuli (Estimate = -0.082, SE = 0.008, z = -9.749, $p < .001$; both real and sham conditions).

Devaluated trials revealed reduced accuracy for overtrained stimuli (i.e. less response switches) compared to the training phase, as shown by an interaction between block x amount of training ($F(1, 9404.14) = 25.689$, $p < .001$). That is, changing the habitual answer was indeed more difficult for the overtrained condition, in line with our hypothesis. For the RT in devaluated trials, participants had higher RT for the response switches in overtrained stimuli compared to the training phase (interaction block x amount of training, ($F(1, 37.691) = 37.691$, $p < .001$)). That is, changing the habitual answer was accompanied by greater cognitive costs in terms of RT for the overtrained condition, which is also in line with our hypothesis. The results align with our hypothesis that there exists more RT switch cost for extended training stimuli when they are devalued.

**Supplementary Figures**

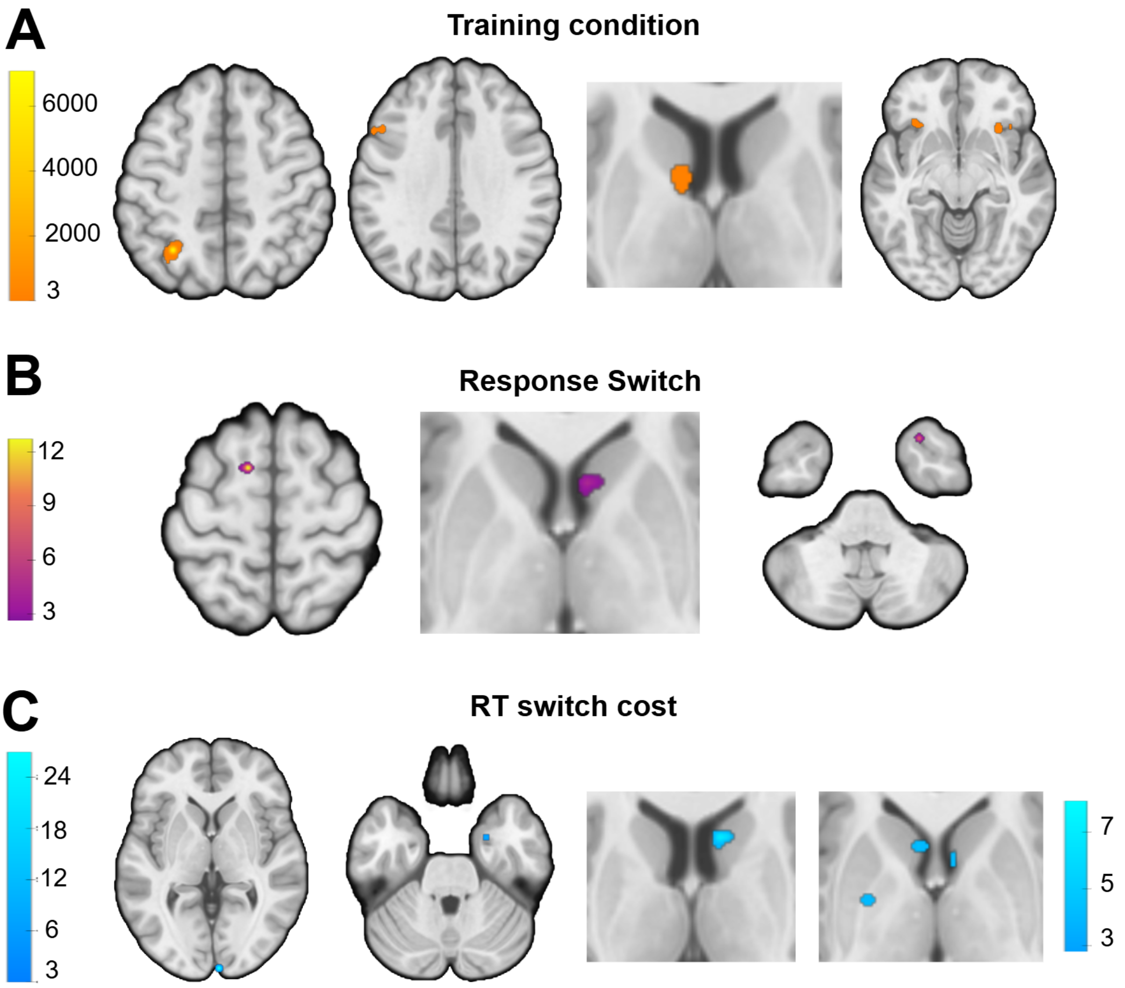


***Figure S1. Brain activity during training and devaluation in goal-directed behaviour (standard training vs overtrained)****. A Bayes factor >= 3 was used for thresholding (as seen in the color bars).* ***A)*** *GLM univariate contrasts showing goal-directed activation during training;* ***B)*** *GLM univariate contrasts engaging goal-directed brain areas when switching the overtrained S-R associations, and* ***C)*** *when adding RT switch costs to model the BOLD data as regressor; the striatal activation in the right side used a small volume corrected analysis with a striatal ROI (color scale shown below the main one). Other additional regions are displayed in tables S1-S3B. No cluster extent threshold was applied, except for the first contrast where we had more trials so we removed single-voxel clusters for visual purposes.*

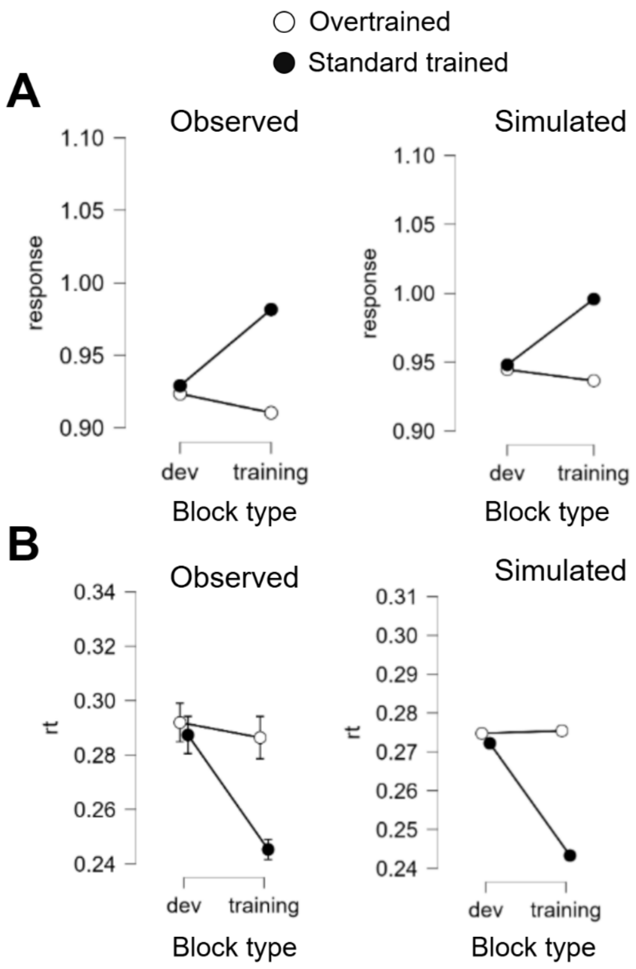


***Figure S2. HDDM posterior predictive check.*** Comparison between simulated and observed behavioral data from the HDDM model in **A)** response accuracy and **B)** reaction times.

**Supplementary tables**

***Table S1: Training phase contrasts (on stimulus time).*** *ROI: small-volume corrected analysis

| **Brain areas** | **Peak value** | **Peak coordinates x, y, z** |
|---|---|---|
| **A. Overtrained > standard trained correct** | | |
| Planum Temporale | 266.5 | 59.5, -24.5, 13.5 |
| Postcentral Gyrus | 34.4 | (23.5, -33.5, 77.5) |
| Superior Parietal Lobule | 9.4 | (17.5, -54.5, 61.5) |
| Postcentral Gyrus | 5.8 | (-27.5, -39.5, 57.5) |
| Precentral Gyrus | 48.4 | (-57.5, -3.5, 33.5) |
| Precentral Gyrus | 7.9 | (-48.5, -78.5, 21.5) |
| Superior Temporal Gyrus posterior division | 11.9 | (-66.5, -45.5, 13.5) |
| Precuneous Cortex | 5.7 | (-9.5, -54.5, 9.5) |
| Lingual Gyrus | 6.6 | (11.5, -66.5, -2.5) |
| Frontal Pole | 5.8 | (-27.5, 44.5, 33.5) |
| Central Opercular Cortex | 5.5 | (44.5, -12.5, 13.5) |
| Cingulate Gyrus posterior division | 4.8 | (17.5, -48.5, 5.5) |
| Insula | 4.6 | (-33.5, -12.5, 17.5) |
| Supramarginal Gyrus anterior division | 3.3 | (65.5, -24.5, 21.5) |
| Supramarginal Gyrus posterior division | 8.9 | (-51.5, -45.5, 37.5) |
| Central Opercular Cortex | 8.5 | (47.5, -12.5, 9.5) |
| Amygdala and hippocampus | 6.3 | (20.5, -6.5, -18.5) |
| Frontal Medial Cortex | 4.6 | (-3.5, 44.5, -22.5) |
| Putamen | 4.2 | (26.5, -9.5, 1.5) |
| Postcentral Gyrus | 4.1 | (-15.5, -39.5, 49.5) |
| Insula | 3.8 | (38.5, -15.5, -2.5) |
| Frontal Medial Cortex | 3.7 | (2.5, 47.5, -22.5) |
| Cerebelum Crus2 R (from AAL atlas) | 3.4 | (53.5, -63.5, -42.5) |
| Precuneous Cortex | 3.3 | (-12.5, -54.5, 13.5) |
| Precentral Gyrus | 3.3 | (-12.5, -30.5, 69.5) |
| Precentral Gyrus; | 3.2 | (50.5, -9.5, 49.5) |
| Cingulate Gyrus anterior division | 3 | (5.5, -9.5, 45.5) |
| *** Striatal ROI:** | | |
| Putamen | 18.6 | (26.5, -9.5, 1.5) |
| Putamen | 4.8 | (-27.5, -15.5, 1.5) |
| Caudate | 3.4 | (14.5, 5.5, 21.5) |
| **B. Standard trained > overtrained correct** | | |
| Superior Parietal Lobule | 10935.9 | (-30.5, -57.5, 49.5) |
| Orbitofrontal cortex | 88 | (-33.5, 26.5, -10.5) |
| Insula | 394.9 | (29.5, 20.5, -6.5) |
| Paracingulate Gyrus | 19.4 | (2.5, 23.5, 37.5) |
| Temporal Occipital Fusiform Cortex | 51.8 | (-42.5, -48.5, -22.5) |
| Middle Frontal Gyrus | 101 | (-48.5, 17.5, 33.5) |
| Precentral Gyrus | 7.8 | (-39.5, 2.5, 33.5) |
| Lateral Occipital Cortex inferior division | 32.9 | (32.5, -84.5, -14.5) |
| Middle Frontal Gyrus | 18.8 | (32.5, 11.5, 29.5) |

| | | |
|---|---|---|
| Caudate | 185.2 | (-9.5, 5.5, 9.5) |
| Middle Frontal Gyrus | 6.3 | (50.5, 23.5, 29.5) |
| Lateral Occipital Cortex superior division | 15.2 | (26.5, -66.5, 45.5) |
| Cingulate Gyrus anterior division | 5.9 | (-3.5, 35.5, 17.5) |
| Superior Temporal Gyrus posterior division | 59.4 | (47.5, -27.5, -2.5) |
| Precuneous Cortex | 5.5 | (8.5, -72.5, 41.5) |
| Cerebelum 8 R | 8.6 | (41.5, -57.5, -50.5) |
| Lateral Occipital Cortex superior division | 8.4 | (32.5, -57.5, 49.5) |
| Lateral Occipital Cortex superior division | 3.8 | (-30.5, -75.5, 25.5) |
| Temporal Occipital Fusiform Cortex | 4.7 | (32.5, -48.5, -22.5) |
| Middle Temporal Gyrus posterior division | 4.7 | (50.5, -15.5, -10.5) |
| Lateral Occipital Cortex superior division | 4.3 | (-24.5, -66.5, 41.5) |
| Lateral Occipital Cortex inferior division; | 4.2 | (-42.5, -72.5, -10.5) |
| Middle Frontal Gyrus | 4.2 | (-45.5, 26.5, 37.5) |
| Occipital Pole | 4.1 | (-3.5, -96.5, -2.5) |
| Cingulate Gyrus anterior division | 3.5 | (-0.5, 8.5, 33.5) |
| Inferior Temporal Gyrus temporooccipital part | 3.2 | (-54.5, -45.5, -14.5) |
| Cerebelum Crus1 L (from AAL atlas) | 3.2 | (-39.5, -57.5, -26.5) |
| *** Striatal ROI** | | |
| Caudate | 1052 | (-9.5, 5.5, 9.5) |
| Caudate | 9.9 | (11.5, 17.5, 1.5) |
| Putamen | 6.8 | (29.5, -0.5, 9.5) |
| Thalamus | 3.1 | (14.5, -0.5, 13.5) |

***Table S2: Response switch contrast (on stimulus time).*** *ROI: small-volume corrected analysis.

| **Brain areas** | **Peak value** | **Peak coordinates x, y, z** |
|---|---|---|
| **A. Response switch overtrained > standard trained** | | |
| Postcentral Gyrus | 19.5 | (-48.5, -15.5, 61.5) |
| Anterior Cingulate Cortex | 456.8 | (5.5, -6.5, 45.5) |
| Cerebellum (6 R) | 64.1 | (29.5, -48.5, -26.5) |
| Central Opercular Cortex | 3.8 | (-51.5, 5.5, -2.5) |
| Supplementary motor area | 10.9 | (-9.5, 2.5, 45.5) |
| Paracingulate Gyrus | 3.9 | (11.5, 44.5, 13.5) |
| *** Striatal ROI:** | | |
| Caudate | 3.1 | (-15.5, 26.5, -2.5) |
| **B. Response switch standard > overtrained** | | |
| Superior Frontal Gyrus | 19 | (-12.5, 5.5, 61.5) |
| Caudate | 6 | (5.5, 14.5, 1.5) |
| Middle Temporal Gyrus posterior division | 4.9 | (-63.5, -24.5, -10.5) |
| Inferior Temporal Gyrus | 4.4 | (59.5, -54.5, -18.5) |
| Temporal Fusiform Cortex posterior division | 3.5 | (-60.5, -12.5, -26.5) |
| Temporal Pole | 12.4 | (29.5, 14.5, -42.5) |
| Frontal Pole | 4.7 | (20.5, 53.5, 17.5) |

| | | |
|---|---|---|
| Frontal Pole | 3.2 | (-18.5, 47.5, 33.5) |
| *** Striatal ROI:** | | |
| Caudate | 11.7 | (-12.5, 11.5, 9.5) |
| Caudate | 28.8 | (5.5, 14.5, 1.5) |
| Putamen | 5.1 | (-6.5, 2.5, -2.5) |
| Putamen | 4.9 | (-30.5, 2.5, -6.5) |
| Putamen | 3.9 | (26.5, -9.5, 5.5) |

***Table S3: RT switch contrast (on stimulus time).*** *ROI: small-volume corrected analysis.

| **Brain areas** | **Peak value** | **Peak coordinates x, y, z** |
|---|---|---|
| **A. RT switch cost overtrained > standard trained** | | |
| Cerebellum (6 R) | 11.7 | (29.5, -48.5, -26.5) |
| Precentral Gyrus | 8.9 | (-18.5, -18.5, 69.5) |
| Superior Parietal Lobule | 7.9 | (20.5, -54.5, 69.5) |
| Planum Temporale | 4.7 | (-36.5, -33.5, 17.5) |
| Superior Temporal Gyrus | 4.2 | (59.5, -12.5, -6.5) |
| Frontal Pole | 13.9 | (-27.5, 41.5, 33.5) |
| Precuneous Cortex | 10 | (8.5, -57.5, 29.5) |
| Frontal Operculum Cortex | 6.7 | (-39.5, 14.5, 9.5) |
| Cerebellum (8 L) | 6.5 | (-24.5, -48.5, -54.5) |
| Precentral Gyrus | 5.6 | (38.5, -9.5, 65.5) |
| Anterior Cingulate cortex | 5.6 | (5.5, -6.5, 45.5) |
| Orbitofrontal cortex | 4.8 | (-9.5, 29.5, -26.5) |
| **B. RT switch cost standard > overtrained** | | |
| Occipital Pole | 44.3 | (5.5, -99.5, 1.5) |
| Temporal Pole | 4.2 | (29.5, 14.5, -42.5) |
| Parahippocampal Gyrus anterior division | 13.6 | (23.5, -0.5, -30.5) |
| Middle Frontal Gyrus | 5.5 | (-30.5, 26.5, 37.5) |
| Temporal Fusiform Cortex | 4.7 | (-42.5, -30.5, -22.5) |
| Frontal Pole | 3.7 | (35.5, 50.5, -2.5) |
| Cerebellum (Crus2 R) | 3.4 | (47.5, -51.5, -46.5) |
| Occipital Pole | 3.3 | (-33.5, -90.5, -10.5) |
| *** Striatal ROI:** | | |
| Caudate | 9.7 | (11.5, 14.5, 9.5) |
| Putamen | 4.9 | (23.5, 14.5, -10.5) |
| Caudate | 8.7 | (-15.5, -9.5, 21.5) |
| Putamen | 3.8 | (-24.5, -3.5, 1.5) |

***Table S4: g-PPI contrast for training phase contrasts (on stimulus time).***

| **Brain areas** | **Peak value** | **Peak coordinates x, y, z** |
|---|---|---|
| **Overtrained correct > Standard trained correct – Seed: S1 (R)** | | |
| Insula | 12.9 | (-30.5, 11.5, -10.5) |
| Lateral Occipital Cortex superior division | 54.4 | (-48.5, -72.5, 29.5) |
| Orbitofrontal cortex | 12 | (-27.5, 29.5, -18.5) |
| Middle Frontal Gyrus | 11.5 | (-33.5, 14.5, 53.5) |
| Subcallosal Cortex | 5.6 | (-12.5, 17.5, -10.5) |
| Superior Frontal Gyrus | 3.2 | (8.5, 5.5, 73.5) |
| Superior Frontal Gyrus | 27.3 | (17.5, 17.5, 61.5) |
| Frontal Pole | 4.5 | (-15.5, 68.5, 17.5) |
| Temporal Pole | 4.4 | (-39.5, 23.5, -34.5) |
| Hippocampus | 4 | (26.5, -12.5, -26.5) |
| Inferior Frontal Gyrus pars triangularis | 3.8 | (-54.5, 23.5, 21.5) |
| **Overtrained correct > Standard trained correct – Seed: Planum temporale (L)** | | |
| Supramarginal Gyrus anterior division | 26.4 | (-54.5, -24.5, 25.5) |
| Lateral Occipital Cortex superior division | 10.6 | (-54.5, -63.5, 17.5) |
| Temporal Occipital Fusiform Cortex | 7 | (38.5, -54.5, -14.5) |
| Precuneous Cortex | 6.3 | (-3.5, -54.5, 5.5) |
| Cerebellum (8 R) | 5.5 | (38.5, -57.5, -54.5) |
| Subcallosal Cortex | 3.1 | (-9.5, 14.5, -18.5) |

***Table S5: g-PPI contrast response switch contrasts (on stimulus time).***

| **Brain areas** | **Peak value** | **Peak coordinates x, y, z** |
|---|---|---|
| **A. Response switch overtrained>Response switch standard trained – Seed: S1 (L)** | | |
| Cuneus | 9.3 | (5.5, -84.5, 29.5) |
| Temporal Pole | 7.7 | (44.5, 8.5, -10.5) |
| Middle Temporal Gyrus posterior division | 5 | (50.5, -3.5, -18.5) |
| Occipital Fusiform Gyrus | 6.2 | (-27.5, -81.5, -6.5) |
| Temporal Pole | 5.8 | (50.5, 17.5, -26.5) |
| Lateral Occipital Cortex superior division | 25.9 | (26.5, -81.5, 17.5) |
| Intracalcarine Cortex | 8.3 | (-6.5, -72.5, 13.5) |
| Temporal Pole | 4.5 | (38.5, 11.5, -18.5) |
| Lingual Gyrus | 3.8 | (-12.5, -72.5, 1.5) |
| Cuneus | 3.2 | (-0.5, -75.5, 25.5) |
| **B. Response switch overtrained > Response switch standard trained – Seed: OFC (L)** | | |
| Inferior Frontal Gyrus pars triangularis | 12.7 | (59.5, 26.5, 9.5) |
| Lateral Ventrical | 9.1 | (-18.5, -39.5, 21.5) |
| Vermis 6 | 8.6 | (2.5, -63.5, -22.5) |
| Inferior Temporal Gyrus | 5 | (53.5, -48.5, -22.5) |
| Lateral Occipital Cortex superior division | 4.2 | (23.5, -78.5, 53.5) |
| Frontal Medial Cortex | 3.5 | (2.5, 53.5, -22.5) |

| | | |
|---|---|---|
| Supplementary motor area | 3.4 | (11.5, -0.5, 53.5) |
| Orbitofrontal cortex | 3.1 | (50.5, 23.5, -10.5) |
| **C. Response switch overtrained > Response switch standard trained – Seed: M1 (middle)** | | |
| Middle Temporal Gyrus temporooccipital part | 8.4 | (50.5, -39.5, -2.5) |
| Parahippocampal Gyrus posterior division | 6.2 | (26.5, -21.5, -22.5) |
| Temporal Occipital Fusiform Cortex | 3.8 | (-30.5, -57.5, -10.5) |
| Superior Temporal Gyrus anterior division | 6.4 | (56.5, -3.5, -6.5) |
| Lateral Occipital Cortex superior division | 3.4 | (29.5, -66.5, 49.5) |
| Occipital Pole | 3.1 | (14.5, -99.5, 9.5) |

***Table S6: g-PPI contrast RT switch cost contrasts (on stimulus time).***

| **Brain areas** | **Peak value** | **Peak coordinates x, y, z** |
|---|---|---|
| **A. RT switch cost overtrained>RT switch cost standard trained – Seed: PMC (L)** | | |
| Precuneous Cortex | 3.4 | (-12.5, -69.5, 21.5) |
| Planum Polare | 31.1 | (-39.5, -0.5, -22.5) |
| Temporal Pole | 22.7 | (-36.5, 20.5, -34.5) |
| Superior Frontal Gyrus | 7.5 | (-24.5, -6.5, 73.5) |
| Supramarginal Gyrus anterior division | 5.9 | (-60.5, -27.5, 21.5) |
| Temporal Pole | 132.5 | (38.5, 11.5, -26.5) |
| Middle Frontal Gyrus | 23 | (35.5, 8.5, 33.5) |
| Precuneous Cortex | 12.7 | (8.5, -51.5, 69.5) |
| Insula | 6.6 | (35.5, -15.5, 9.5) |
| Heschl's Gyrus (includes H1 and H2) | 6.3 | (38.5, -18.5, 9.5) |
| Hippocampus | 5.2 | (-30.5, -36.5, -6.5) |
| Pallidum | 4.9 | (17.5, 5.5, -2.5) |
| Putamen | 4.7 | (-30.5, -9.5, 1.5) |
| Temporal Fusiform Cortex posterior division | 3.6 | (-33.5, -42.5, -18.5) |
| Precentral Gyrus | 3.2 | (32.5, -15.5, 69.5) |
| **B. RT switch cost overtrained > RT switch cost standard trained – Seed: S2 (R)** | | |
| Precentral Gyrus | 7.3 | (-6.5, -15.5, 61.5) |
| Angular Gyrus | 4.7 | (59.5, 5.5, -22.5) |
| Insula | 5.2 | (-33.5, -18.5, 5.5) |
| Middle Temporal Gyrus posterior division | 5 | (62.5, -18.5, -18.5) |
| Precuneous Cortex | 4.2 | (2.5, -54.5, 9.5) |
| Frontal Medial Cortex | 3.7 | (-0.5, 32.5, -14.5) |
| Middle Temporal Gyrus posterior division | 3.1 | (-69.5, -39.5, -2.5) |

***Table S7. DIC values of the different model variations tested for the HDDM.*** Higher values indicates better fits for each model.

| **Model** | **DIC** |
|---|---|
| Complete unified model | -9054.270270 |
| Separating conditions for "t" | -8656.318142 |
| Separating conditions for "a" | -8517.505693 |
| Separating conditions for "v" | -8646.397243 |
| Fixing t=0 | -8783.431136 |

***Table S8: Outcome condition contrasts (on feedback time).*** *ROI: small-volume corrected analysis.

| **Brain areas** | **Peak value** | **Peak coordinates x, y, z** |
|---|---|---|
| **A. Overtrained > standard trained correct** | | |
| Posterior Cingulate Cortex | 284.5 | (-12.5, -48.5, 1.5) |
| Central Opercular Cortex | 17.6 | (62.5, -12.5, 9.5) |
| Precuneous | 11.0 | (-9.5, -57.5, 21.5) |
| Postcentral Gyrus | 9.9 | (-24.5, -18.5, 73.5) |
| Fronto-medial cortex | 17.3 | (-9.5, 41.5, -10.5) |
| Precuneous | 6.4 | (17.5, -57.5, 17.5) |
| Thalamus | 6.6 | (14.5, -24.5, 9.5) |
| Temporal Occipital Cortex | 6.2 | (32.5, -45.5, -6.5) |
| Amygdala | 44.5 | (-15.5, -3.5, -14.5) |
| Middle Temporal Gyrus | 12.2 | (-57.5, -0.5, -18.5) |
| Lateral Occipital Cortex | 6.4 | (29.5, -90.5, 25.5) |
| Frontal Medial Cortex | 4.0 | (5.5, 50.5, -10.5) |
| Amygdala | 7.2 | (20.5, -6.5, -18.5) |
| Cingulate Cortex | 6.1 | (11.5, 41.5, 13.5) |
| Hippocampus | 4.4 | (29.5, -33.5, -10.5) |
| Orbito Frontal Cortex | 5.2 | (-12.5, 8.5, -14.5) |
| *** Striatal ROI:** | | |
| Nucleus Accumbens | 70.5093 | -9.5, 5.5, -14.5 |
| Nucleus Accumbens | 7.52511 | 11.5, 8.5, -10.5 |
| Putamen | 8.49437 | -18.5, 8.5, -6.5 |
| Putamen | 5.43378 | 26.5, -.65, 1.5 |
| Caudate | 10.0306 | -15.5, 2.5, 13.5 |
| **B. Standard > overtrained trained correct** | | |
| Superior Parietal Lobule | 81.2 | (-45.5, -51.5, 61.5) |
| Occipital Pole | 65.3 | (-9.5, -93.5, -18.5) |
| Lateral Occipital Cortex | 11.4 | (-36.5, -60.5, 61.5) |
| Cerebelum (Crus1) | 28.8 | (-21.5, -84.5, -22.5) |
| Cerebelum (Crus2) | 8.4 | (-15.5, -75.5, -38.5) |
| Lateral Occipital Cortex | 14.5 | (29.5, -81.5, -22.5) |
| Orbito Frontal Cortex | 4.4 | (29.5, 23.5, -6.5) |

*** Striatal ROI:**

| | | |
|---|---|---|
| Caudate_R | 6.9303 | 17.5, -18.5, 21.5 |
| Putamen_R | 6.27586 | 26.5, -3.5, 9.5 |
| Putamen_L | 4.5244 | -21.5, 8.5, 1.5 |

Gorgolewski, Krzysztof, Christopher D. Burns, Cindee Madison, Dav Clark, Yaroslav O. Halchenko, Michael L. Waskom, and Satrajit S. Ghosh. 2011. "Nipype: A Flexible, Lightweight and Extensible Neuroimaging Data Processing Framework in Python." *Frontiers in Neuroinformatics* 5 (August): 13.

Abraham, Alexandre, Fabian Pedregosa, Michael Eickenberg, Philippe Gervais, Andreas Mueller, Jean Kossaifi, Alexandre Gramfort, Bertrand Thirion, and Gaël Varoquaux. 2014. "Machine Learning for Neuroimaging with Scikit-Learn." *Frontiers in Neuroinformatics* 8 (February): 14.

Avants, B. B., C. L. Epstein, M. Grossman, and J. C. Gee. 2008. "Symmetric Diffeomorphic Image Registration with Cross-Correlation: Evaluating Automated Labeling of Elderly and Neurodegenerative Brain." *Medical Image Analysis* 12 (1): 26–41.

Behzadi, Yashar, Khaled Restom, Joy Liau, and Thomas T. Liu. 2007. "A Component Based Noise Correction Method (CompCor) for BOLD and Perfusion Based FMRI." *NeuroImage* 37 (1): 90–101.

Cox, R. W., and J. S. Hyde. 1997. "Software Tools for Analysis and Visualization of FMRI Data." *NMR in Biomedicine* 10 (4–5): 171–78.


Esteban, Oscar, Christopher J. Markiewicz, Ross W. Blair, Craig A. Moodie, A. Ilkay Isik, Asier Erramuzpe, James D. Kent, et al. 2019. "FMRIPrep: A Robust Preprocessing Pipeline for Functional MRI." *Nature Methods* 16 (1): 111–16.


Fonov, V. S., A. C. Evans, R. C. McKinstry, C. R. Almli, and D. L. Collins. 2009. "Unbiased Nonlinear Average Age-Appropriate Brain Templates from Birth to Adulthood." *NeuroImage* 47 (July): S102.


Gorgolewski, Krzysztof, Christopher D. Burns, Cindee Madison, Dav Clark, Yaroslav O. Halchenko, Michael L. Waskom, and Satrajit S. Ghosh. 2011. "Nipype: A Flexible, Lightweight and Extensible Neuroimaging Data Processing Framework in Python." *Frontiers in Neuroinformatics* 5 (August): 13.


Greve, Douglas N., and Bruce Fischl. 2009. "Accurate and Robust Brain Image Alignment Using Boundary-Based Registration." *NeuroImage* 48 (1): 63–72.

Jenkinson, Mark, Peter Bannister, Michael Brady, and Stephen Smith. 2002. "Improved Optimization for the Robust and Accurate Linear Registration and Motion Correction of Brain Images." *NeuroImage* 17 (2): 825–41.

Lanczos, C. 1964. "Evaluation of Noisy Data." *Journal of the Society for Industrial and Applied Mathematics Series B Numerical Analysis* 1 (1): 76–85.

Power, Jonathan D., Anish Mitra, Timothy O. Laumann, Abraham Z. Snyder, Bradley L. Schlaggar, and Steven E. Petersen. 2014. "Methods to Detect, Characterize, and Remove Motion Artifact in Resting State FMRI." *NeuroImage* 84 (January): 320–41.

Tustison, Nicholas J., Brian B. Avants, Philip A. Cook, Yuanjie Zheng, Alexander Egan, Paul A. Yushkevich, and James C. Gee. 2010. "N4ITK: Improved N3 Bias Correction." *IEEE Transactions on Medical Imaging* 29 (6): 1310–20.

Zhang, Y., M. Brady, and S. Smith. 2001. "Segmentation of Brain MR Images through a Hidden Markov Random Field Model and the Expectation-Maximization Algorithm." *IEEE Transactions on Medical Imaging* 20 (1): 45–57.

Esteban, Oscar, Daniel Birman, Marie Schaer, Oluwasanmi O. Koyejo, Russell A. Poldrack, and Krzysztof J. Gorgolewski. 2017. "MRIQC: Advancing the Automatic Prediction of Image Quality in MRI from Unseen Sites." *PloS One* 12 (9): e0184661.
